# PCNA unloading is crucial for the bypass of DNA lesions by homologous recombination

**DOI:** 10.1101/2024.02.18.580888

**Authors:** Matan Arbel-Groissman, Batia Liefshitz, Nir Katz, Maxim Kuryachiy, Martin Kupiec

## Abstract

The DNA Damage Tolerance (DDT) mechanisms allow cells to bypass lesions in the DNA during replication. This allows the cells progress normally through the cell cycle in face of abnormalities in their DNA. PCNA, a homotrimeric sliding clamp complex plays a central role in the coordination of various processes during DNA replication, including the choice of mechanism used during DNA damage bypass. Mono-or poly-ubiquitination of PCNA facilitate an error-prone or an error-free bypass mechanism, respectively. In contrast, SUMOylation recruits the Srs2 helicase, which prevents local homologous recombination. The Elg1 RFC-like complex plays an important role in unloading PCNA from the chromatin. We analyse the interaction of mutations that destabilize PCNA with mutations in the Elg1 clamp unloader and the Srs2 helicase. Our results suggest that in addition to its role as a coordinator of bypass mechanisms, the very presence of PCNA on the chromatin prevents homologous recombination, even in the absence of the Srs2 helicase. Thus, PCNA unloading seems to be a pre-requisite for recombinational repair.

## Introduction

During the cell cycle, the DNA faces numerous attacks, from exogenous or endogenous sources. Damaged DNA, if left untreated, can cause stalling or even collapse of the replication fork. A variety of DNA repair mechanisms exists in order to repair the damaged DNA, and additional mechanisms are able to bypass lesions during DNA replication, allowing their repair in a post-replicative manner [1]. Many of the DNA repair and damage bypass mechanisms, which can collectively be called DNA Damage Tolerance (DDT) pathways are orchestrated by post-translational modifications of the PCNA clamp [2,3]. PCNA is a homotrimeric complex, composed in yeast of three identical Pol30 proteins, which serves as a sliding clamp and processivity factor for the different DNA polymerases [4]. PCNA is modified by a variety of post-translational modifications (PTMs), including ubiquitination, SUMOylation, and phosphorylation. These PTMs regulate the activity of PCNA and its interaction with many proteins involved in DNA replication and DDT; most PTMs are tightly evolutionarily conserved [5].

In response to DNA damage, PCNA is modified by ubiquitination at lysine 164 [6]. This ubiquitination, carried out by the Rad6-Rad18 complex, signals for the recruitment of DNA damage tolerance (DDT) proteins to the damaged site. Whereas mono-ubiquitination activates an error-prone DNA damage bypass orchestrated in yeast by Pol-Zeta [7,8], further poly-ubiquitination of PCNA by Rad5-Mms2-Ubc13 facilitates a template switch bypass of the lesion, in an error-free manner [9–11].

PCNA is loaded on the chromatin by the replication factor C complex (RFC) composed of the Rfc1-5 proteins, and is unloaded by a complex which shares with RFC the four small subunits Rfc2-5, but contains Elg1 instead of Rfc1 (the Elg1 RFC-like complex or RLC) [12]. The timely unloading of PCNA by the Elg1 RLC plays a major role in replication and DNA damage bypass and repair [13,14]. Defects in Elg1 (ATAD5 in mammals) function lead to increased chromosome instability, elevated recombination and mutation rates, and additional genomic instability phenotypes [15]. In the absence of Elg1, PCNA, and in particular SUMOylated PCNA accumulate on the chromatin [16], suggesting that SUMOylation of PCNA may be a signal for its recruitment [17]. PCNA undergoes SUMOylation at lysines 164 and 127 [3].

Elg1 has a complicated genetic interaction with Srs2, another protein that is recruited to PCNA following its SUMOylation [18,19]. Srs2 is a DNA helicase that plays a critical role in the regulation of DNA damage repair and bypass pathways in budding yeast [20–22]. Following DNA damage, ssDNA may be created, which is bound by RPA, and later replaced by Rad51, forming a nucleoprotein filament [23]. This filament can interfere with DNA replication and transcription [24]. The Srs2 helicase unwinds the Rad51 filament and removes it from the DNA. This activity is necessary during the repair of double-stranded breaks [25], but also prevents the formation of unwanted recombination intermediates and allows the cell to repair the DNA damage by other mechanisms. [26,27] Srs2 is recruited to PCNA following its SUMOylation at lysines 164 or 127 [21]. By evicting Rad51 from the DNA, it down- regulates the alternative, homologous recombination-based “salvage pathway”[28].

In this paper we show that PCNA plays a role not only in recruiting and coordinating DNA damage repair proteins, but also that its presence on DNA, even in the absence of Srs2, can serve to prevent recombination in certain situations. By enclosing DNA, PCNA may sometimes serve as a physical constraint against recombinational repair.

## Results

### Spontaneous unloading of PCNA supresses the sensitivity conferred by an inactive DDT mechanism

Because Pol30 is an essential protein, it is not possible to study its function by deleting the gene that encodes it. To overcome this limitation, we made use of Pol30 mutants that form a PCNA complex that is prone to spontaneously disassemble and fall from the chromatin [29,30]. This allows us to measure how the cells react to low levels of PCNA on the chromatin. Numerous such DPP (<u>D</u>isassembly <u>P</u>rone <u>P</u>CNA) mutants were characterized. We focused on two mutations, E143K and V180D, which were shown to lower the affinity of the Pol30 subunits to each other. Without showing differences in the level of expression of the *POL30* gene, *pol30-E143K* mutants were shown in the past to exhibit a mild, and *pol30-V180D* a strong, reduction in the level of PCNA on the chromatin [29]. We confirmed these results in our genetic background (Figure 1).

**Figure 1:**
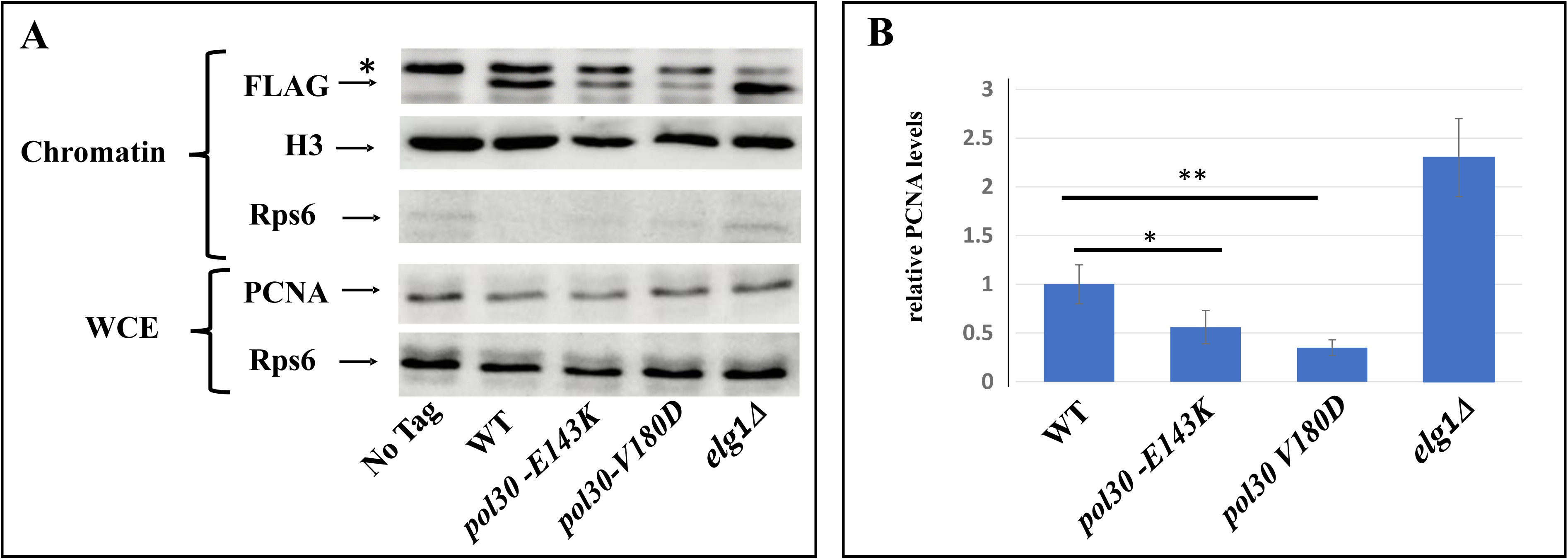
**A.** The upper three panels are Western blots of the chromatin fraction showing flag-tagged PCNA levels on the chromatin, with histone H3 as a nuclear marker for quantification and Rps6 as a cytoplasmic marker. The lower two panels show PCNA and RPS6 from the whole cell extract (WCE) as a control. **B.** Quantitation of this experiment together with 3 additional replicates.

Lysine 164 of Pol30 is the main residue of PCNA that undergoes ubiquitination and SUMOylation [7]. Ubiquitination of K164 is a prerequisite for the activation of most of the DDT pathways [28], and as such, mutations that replace this lysine greatly sensitize the cells to DNA damaging agents as both the error prone and error free repair pathways are inactive [7]. It was previously noted that, perhaps counter-intuitively, an additional mutation in residue K127 supresses the sensitive K164R phenotype [7]. This is due to the fact that SUMOylation of lysine 127 can still recruit Srs2, even in a strain mutated for lysine 164. By evicting Rad51 from the DNA, Srs2 down-regulates the salvage pathway (which is functional in the single *pol30-K164R* mutant). In a double mutant (*pol30-K127R,K164R*) Srs2 is not recruited, and the salvage pathway remains open, providing repair to the damaged DNA and reducing the sensitivity [19].

Surprisingly, the DPP mutations are able to supress the DNA damage sensitivity of *pol30-K164R* strains (Figure 2). Adding the *pol30-V180D* mutation restores the *pol30-K164R* mutant to the same level of sensitivity conferred by the *pol30-K127R* mutation, whereas the suppression by *pol30-E143K* is slightly weaker, but clear (Figure 2). Given the stronger effect of *pol30-V180D,* we continued characterizing this mutation in particular.

**Figure 2:**
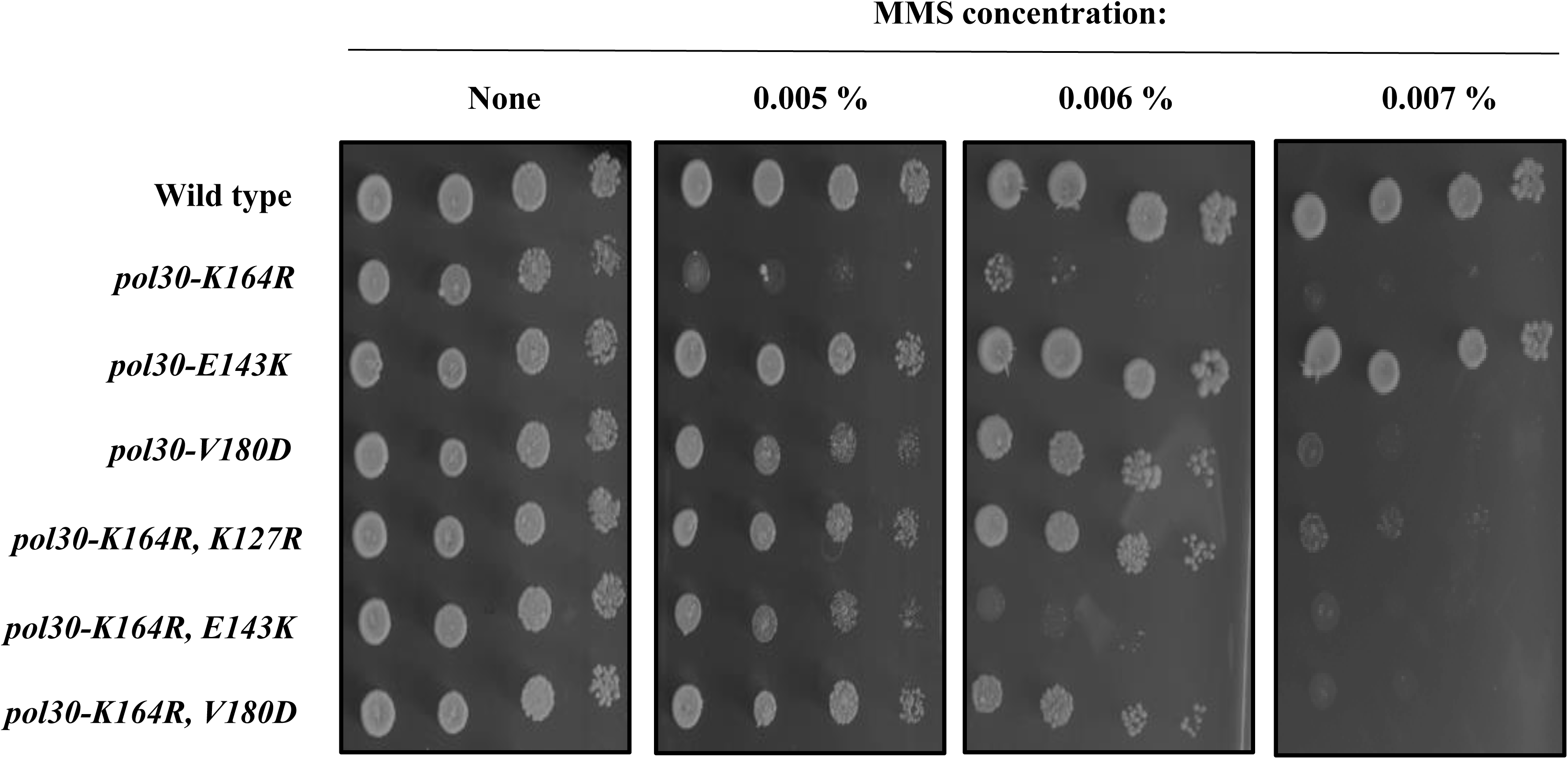
**A.** Ten-fold dilution spot assay on plates with increasing amounts of methylmethane sulfonate (MMS). *pol30-V180D* confers mild sensitivity to DNA damage, to the same extent as *pol30-K164R, K127R*, while *pol30-E143K* does not cause any apparent DNA damage sensitivity. The *pol30-K164R* mutation greatly sensitizes the cells to DNA damage, and is suppressed by an additional mutation, either *pol30-V180D* or *pol30-K127R*, and to a lesser extent, *pol30-E143K*.

### Unstable PCNA results in low amount of Srs2 on the chromatin

The most logical explanation to the fact DPP mutations supress the sensitivity of *pol30-K164R* is that the reduced amount of PCNA on the chromatin de-facto lowered the amount of Srs2 on the chromatin, thus activating the salvage pathway. The hypothesis arising from this model is that the suppression will be similar in level to that provided by *pol30-K127R* and by deletion of *SRS2* and will be epistatic to both. Figure 3 shows that indeed this is the case. The *pol30-K164R,K127R* and *pol30-K164R,V180D*, and even the triple *pol30-K164R, K127R,V180D* mutant, show the same sensitivity to MMS whether the Srs2 helicase is active or completely absent (Figure 3).

**Figure 3:**
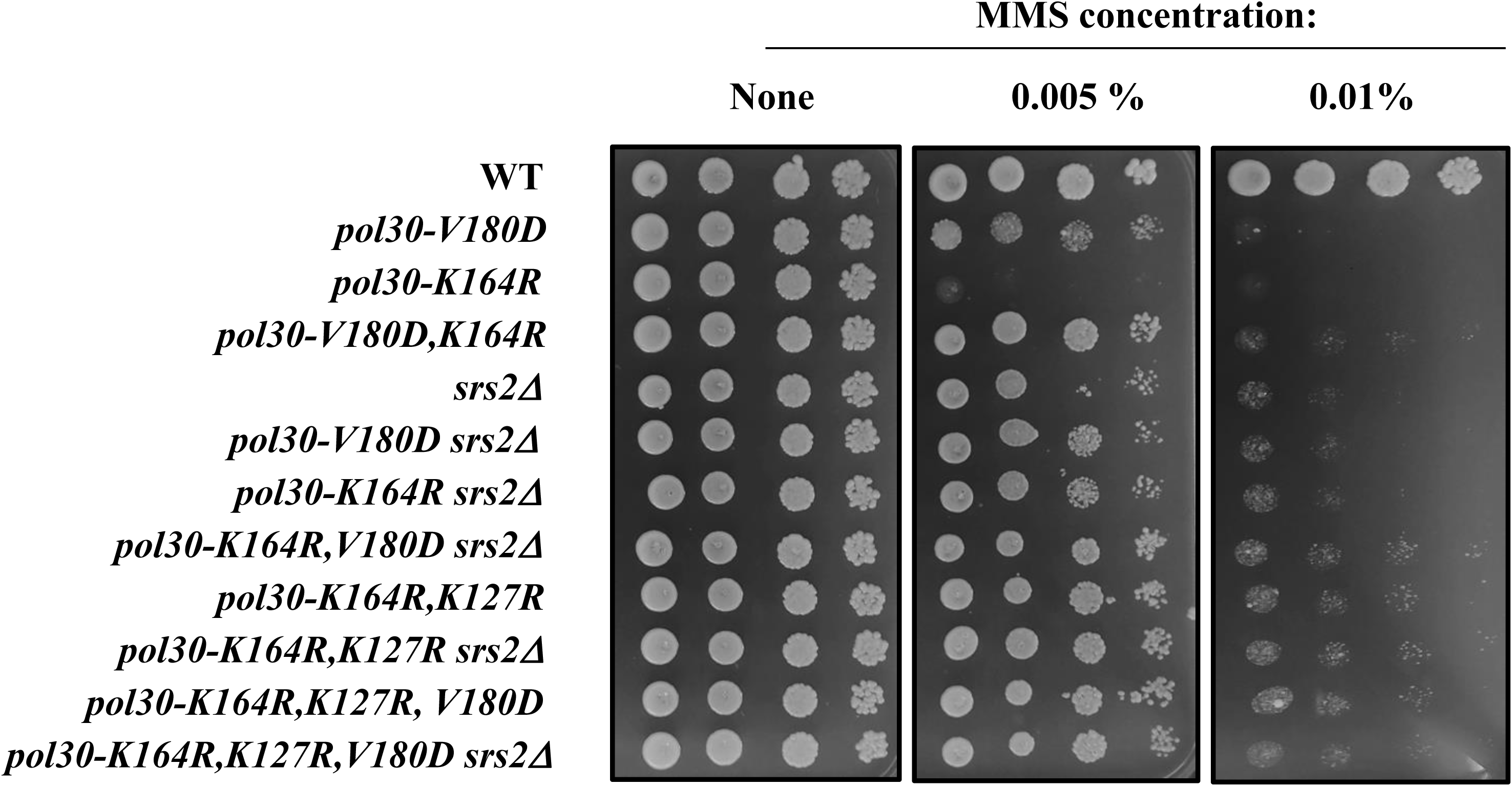
Ten-fold dilution spot assay on plates with increasing amounts of methylmethane sulfonate (MMS). The sensitivity of *pol30-K164R* is suppressed to the same extent by deleting *SRS2,* by introducing the DPP mutations *pol30-K127R* or *pol30-V180D* or by their combinations.

To further validate this claim, we checked the level of Srs2 protein on the chromatin in the different mutants (Figure 4). As expected, mutating lysine 164 of PCNA decreases the amount of Srs2 on the chromatin, consistent with the role of lysine 164 as the main SUMOylation site of the Pol30 protein. The DPP mutation *pol30-V180D* on its own reduces Srs2 levels on the chromatin even more than *pol30-K164R*. Importantly, when the two mutations are combined, the levels of Srs2 on the chromatin are almost undetectable, similarly to what is observed in a *pol30-K164R, K127R* double mutant that lacks PCNA SUMOylation altogether (Figure 4).

**Figure 4:**
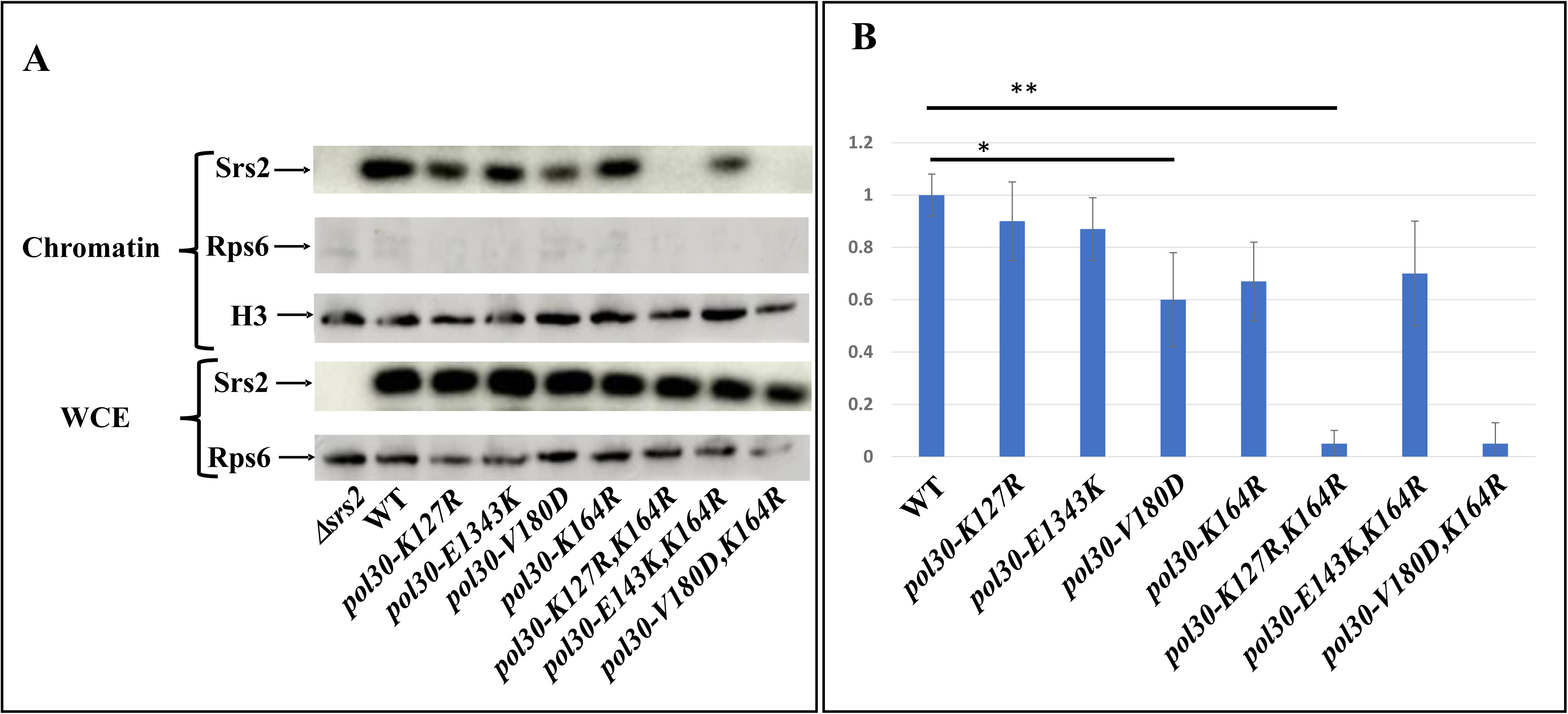
The upper three panels are Western blots of the chromatin fraction showing Srs2 levels on the chromatin, with histone H3 as a nuclear marker for quantification and RPS6 as a cytoplasmic marker. The lower two panels show Srs2 and Rps6 from the whole cell extract (WCE) as a control. **B.** Quantitation of this experiment together with 3 additional replicates.

### DPP suppression of *pol30-K164R* is dependent on homologous recombination (HR)

If indeed, as hinted by the results from the previous section, the DPP mutation activates the salvage pathway by lowering the amount of Srs2 on the chromatin thus, this suppression should be dependent on HR proteins [31]. Most HR reactions in the cell depend on the activity of Rad52 [32,33]. As *rad52*Δ cells are extremely sensitive to MMS, we turned to UV, a DNA damaging agent to which they are relatively resilient, allowing us to check the relationship between the gene deletions. In Figure 5 we show that whereas neither *RAD52* deletion nor any of the PCNA mutations render the strain sensitive to the inflicted UV levels, combining *rad52Δ* with either *pol30-K164R, K127R* or with *pol30-K164R,V180D* greatly sensitizes the cells. We believe that this is because both double mutants are unable to recruit Srs2 to the fork. In the absence of ubiquitination at lysine 164, which prevents opening of the DDT, cells use the salvage pathway, which is now open due to the lack of Srs2 [19]. Without Rad52, however, this mechanism fails, resulting in greatly increased sensitivity to UV. These results demonstrate that, similarly to the suppression of *pol30-K164R* by *pol30-K127R* [21], the suppression by *pol30-V180D* is due to the activity of the salvage pathway. Interestingly, deletion of *RAD52* causes a mild increase in the sensitivity of *pol30-V180D;* this is possibly because the spontaneous unloading of PCNA in this mutant precludes the use of DDT pathways and thus results in an increased dependency on the salvage pathway. At the same time, the difference in sensitivity of the double, when compared to the single *rad52Δ* mutant, implies that PCNA plays an important role in the repair of UV damage when HR is not an option.

**Figure 5:**
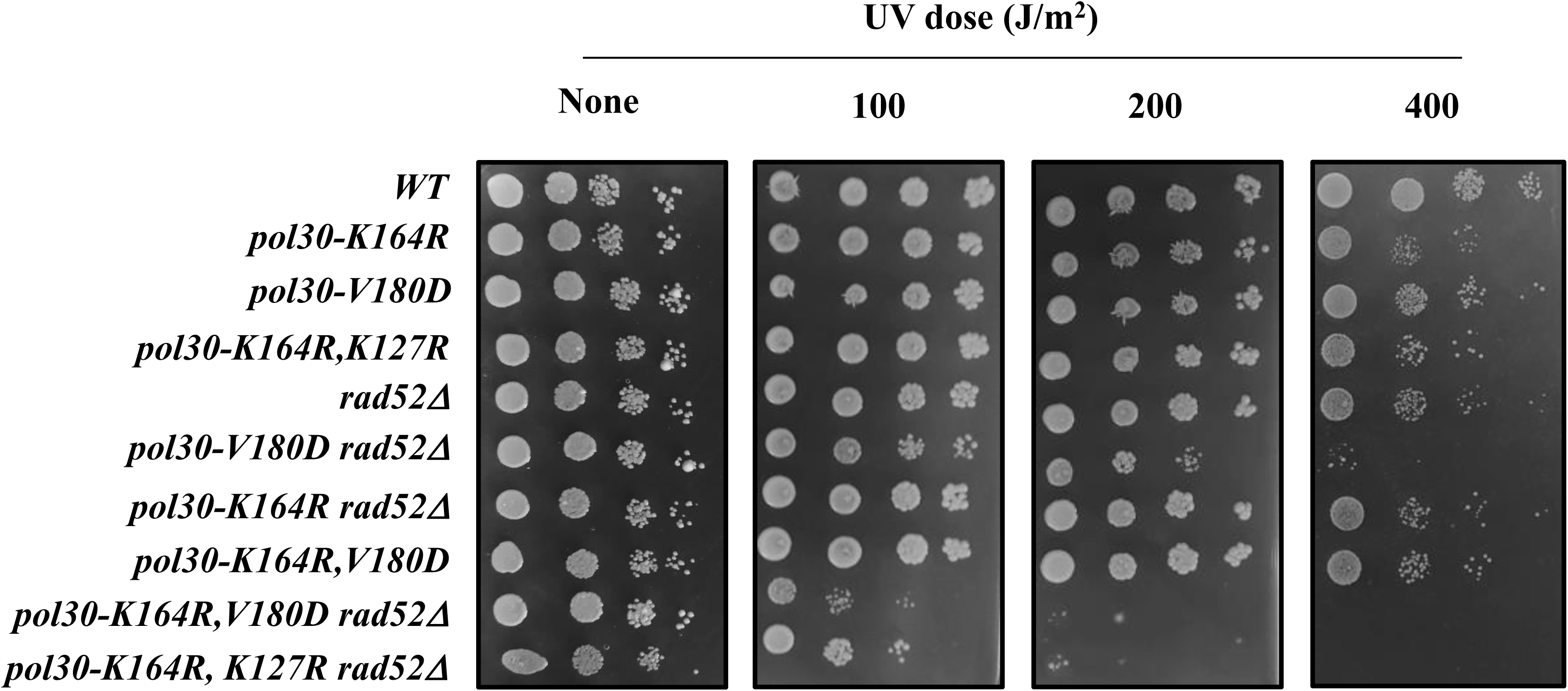
Ten-fold dilution spot assay on plates subjected to increased UV irradiation. The suppression of the DNA damage sensitivity of *pol30-K164R* by *pol30-K127R* and *pol30-V180D* is dependent on the homology recombination factor Rad52.

### Suppression of the synthetic sickness of *elg1Δsrs2Δ* double mutants by spontaneous unloading of PCNA

Whereas singly deleting either the *ELG1* or the *SRS2* genes does not result in a great increase in DNA damage sensitivity, the *elg1Δ srs2Δ* double mutant is extremely sick and exhibits great sensitivity to DNA damage[34]. This can be due to the fact that PCNA may act as a constraint, preventing HR. In the absence of Srs2, the salvage pathway may be activated, but the increased levels of PCNA on chromatin (due to lack of clamp unloading in the absence of Elg1) may curtail its performance. Figure 6A shows that the extreme DNA damage sensitivity of *elg1Δ srs2Δ* double mutants can be completely rescued by the *pol30-V180D* mutation, implying that spontaneous unloading of PCNA suppresses the DNA damage sensitivity of the strain, thus strengthening our claim. Moreover, the *pol30-V180D* mutation is also able to supress the synthetic sickness of the double mutant strain (Figure 6B), again supporting the claim that PCNA serves as a physical constraint against homologous recombination.

**Figure 6:**
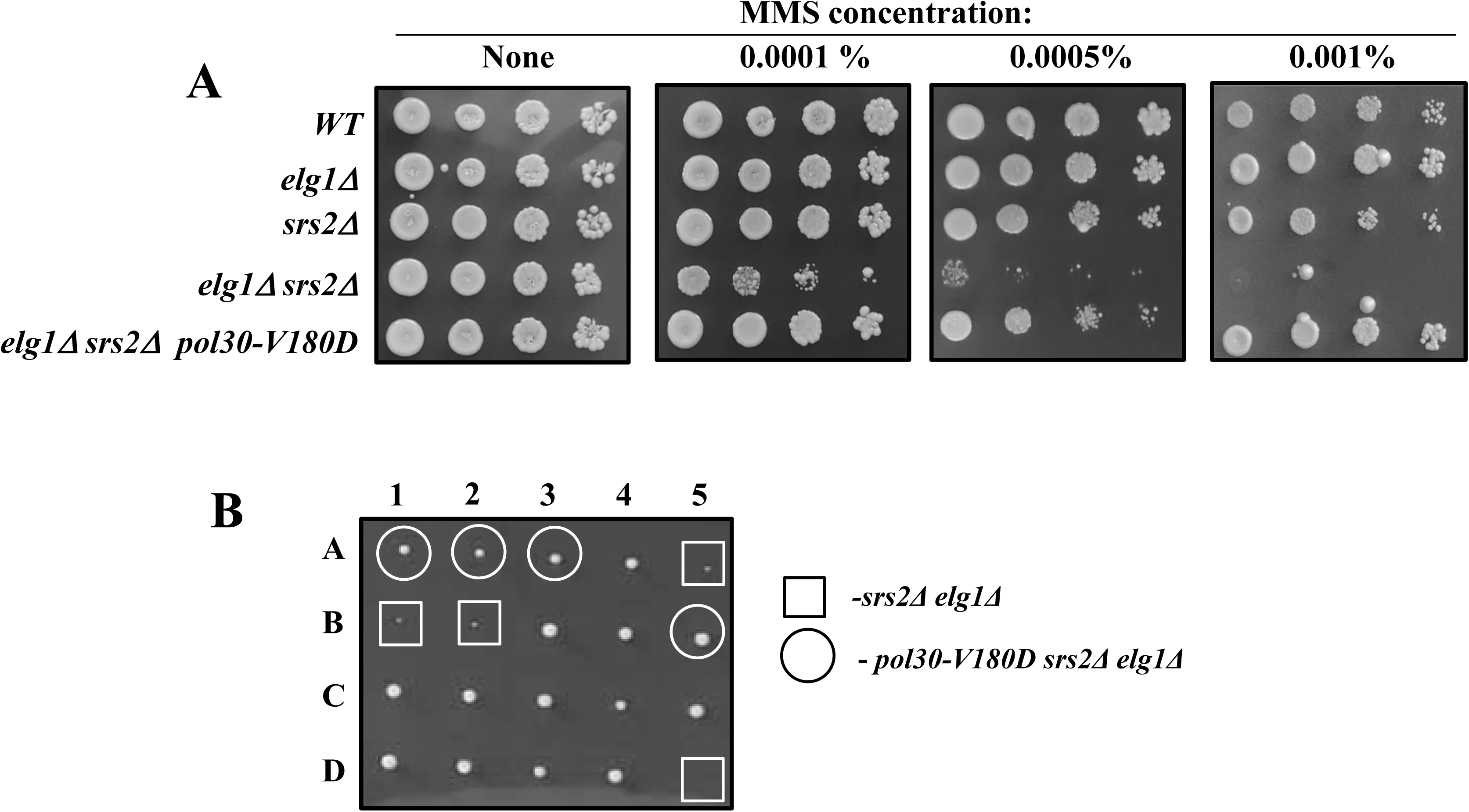
**A.** Ten-fold dilution spot assay on plates with increasing amounts of methylmethane sulfonate (MMS). **B.** Tetrad analysis of a diploid created by mating a *pol30-V180D srs2Δ* haploid with an *elg1Δ* haploid of opposite mating type. Whereas the double mutant *elg1Δ srs2Δ* is sick (squares), the triple mutant (circles) is not.

## Discussion

Despite the important role of the damage bypass mechanisms in cell survival and in the prevention of cancer, as well as their role in evolution, their regulation remains poorly understood. One of the more interesting but puzzling points in the DDT pathways is the role of Srs2 and the activity of the salvage pathway. The presence of Srs2 at the fork, recruited by SUMOylation of PCNA, actively discourages recombination by preventing the formation of Rad51 filaments on DNA [26], a prerequisite for most homologous recombination (HR) events [32]. This, in turn, allows the activation of the DDT pathways, either mutagenic or error free. When Srs2 is absent from the fork the DNA lesion can be bypassed using HR, circumventing the need for any of the other DDT pathways or proteins [35]. PCNA ubiquitination at lysine 164 acts as a master activation switch for the DDT pathways. Therefore mutations that prevent that ubiquitination confer extreme sensitivity to DNA damage [21]. Preventing the recruitment of Srs2 by deleting the gene, or mutating the second lysine that can undergo SUMOylation (K127) partially suppress the sensitivity. Here we have shown that similar results can be obtained by introducing mutations that cause spontaneous disassembly of PCNA, such as *pol30-V180D* or *pol30-K143D*, which reduce the amount of PCNA on chromatin (Figures 2 and 3).

Our results support a model in which the very presence of PCNA on the chromatin constrains the ability of the cells to bypass lesions by using the recombinational salvage pathway. The fact that it is enough to delete *SRS2* to bypass the sensitivity of any other mutation in the DDT pathways implies that when Srs2 is not there, all repair or bypass is canalized towards HR. For example, deletion of any of the genes of the error-free repair pathway group usually results in greatly elevated mutation rate, as now all repair is being send to the error-prone repair pathways, but, when combined with Srs2 deletion, the elevated mutation rate is suppressed [35,36]. This demonstrates again that the lack of Srs2 from the fork results in the uncontrolled activation of the salvage recombination and under these circumstances all the other DDT pathways are unused. So why is the deletion of *ELG1* so harmful to strains lacking Srs2? The most plausible explanation, as previously stated, is that once the process of HR has started, the inability of cells lacking Elg1 to unload PCNA results in recombination intermediates that are left unresolved, leading to cell toxicity and death. Figure 7 shows a schematic model of the bypass of a DNA lesion in Srs2 deficient strain in a post-replicative manner. PCNA timely unloading and reloading is important for allowing the strand exchange, as PCNA encloses a DNA duplex in which one of the strands is damaged, but repair requires an exchange. PCNA should be unloaded and further re-loaded since the bypass requires copying of the sister DNA duplex, and thus the enclosing of the newly synthesized strand and the complementary one from the sister chromatid.

**Figure 7:**
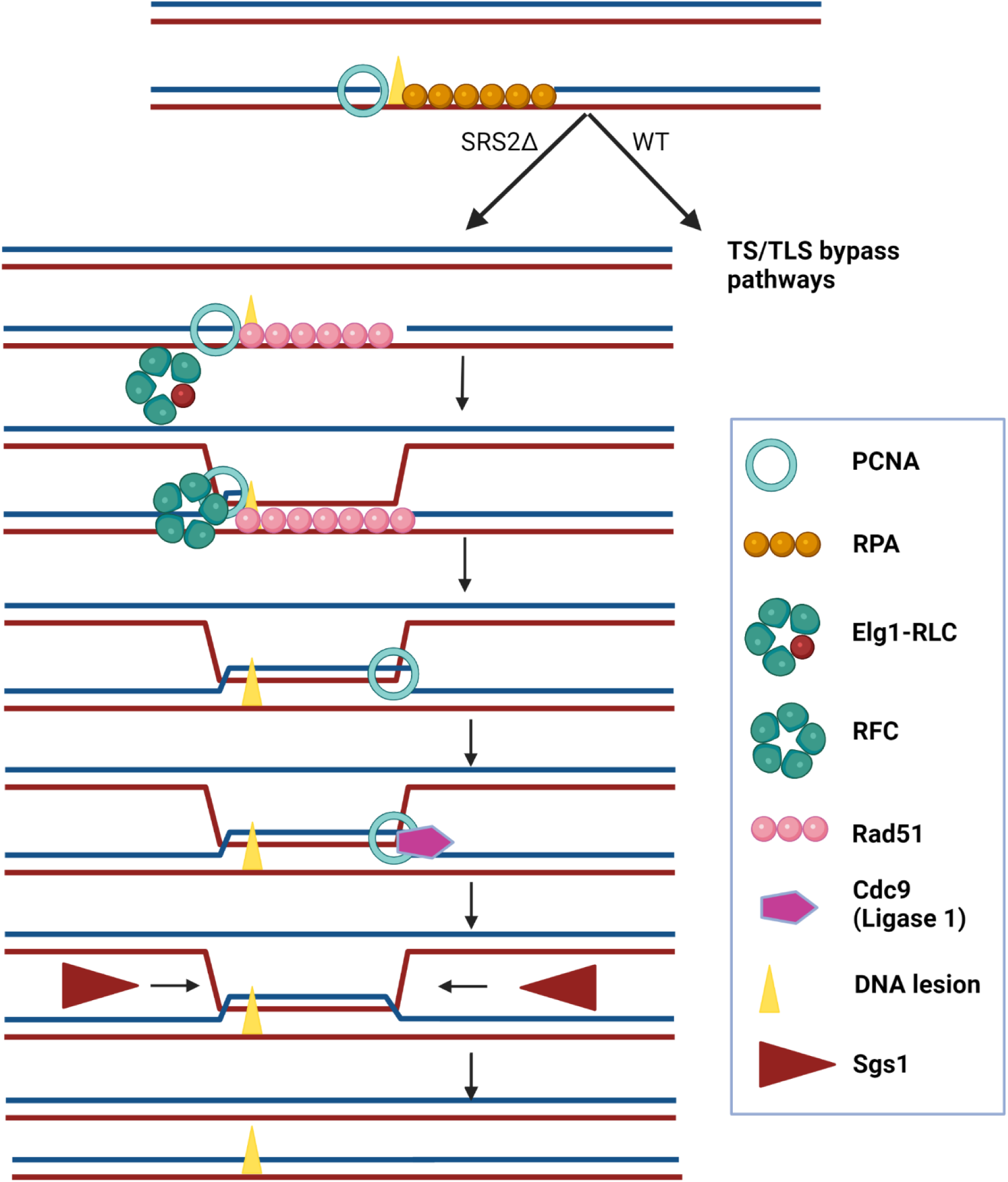
A schematic illustration of the post-replicative bypass of a ssDNA gap left behind the fork in Srs2-deficient cells. An cell in which a replication fork has been arrested due to the presence of a lesion has left a ssDNA gap behind, which is rapidly filled up by RPA. Whereas is w.t. cells ubiquitination of PCNA would promote error-free or error-prone DDT, in the absence of Srs2 RPA is replaced by Rad51, PCNA is unloaded by the Elg1 RLC, and re-loaded (by RFC) to allow copying of the information present in the sister chromatid. After ligation (by Cdc9) the intertwined chromatids are resolved by the Sgs1 helicase and the Top3/Rmi1 topoisomerase activity (not shown).

PCNA unloading importance in salvage recombination pathway explains why a Pol30 mutant that spontaneously disassembles from the chromatin can suppress the synthetic sickness of *elg1Δ srs2*Δ strains. When the salvage pathway is continually activated (as is the case in *srs2Δ*), PCNA has to be timely unloaded or it will lead to fork arrest. An alternative explanation, although less probable in our opinion, is that there is a third partner to this dance: a mysterious protein that binds to PCNA and when mounted on the chromatin causes havoc and damage to the cell. This harmful protein would be normally competing with Srs2 for binding to PCNA and could be usually regulated also by the unloading of PCNA by Elg1. When both Elg1 and Srs2 are deleted, the lack of competition for binding to PCNA and the elevated levels of chromatin bound PCNA results in high amount of the toxic protein on the chromatin, leading to the synthetic sickness and extreme sensitivity. We believe that such a protein would have already caught the attention of the scientific community, although this remains a formal possibility. Future genetic screens may confirm or deny this possibility.

Srs2 and Elg1 have a complicated dance to perform throughout the cell cycle. Whereas Srs2 moves along with the fork during S phase [18] preventing local recombination events, Elg1 is also continuously needed at the fork to cycle PCNA, at each Okazaki fragment during the replication of the lagging strand [37]. They both have SUMO-interacting motifs, and exhibit a preference for SUMOylated PCNA. Several important questions remain. How is Srs2 unloaded? Does the Elg1 RLC normally unload it together with the SUMOylated PCNA? Do Srs2 and Elg1 compete for the binding to SUMOylated PCNA? A future avenue to pursue is understanding the temporal and spatial regulation of Srs2 presence and activity. As it stands now, almost all our knowledge of this pathway is derived from experiments that include an artificial removal of Srs2 from the fork. When, and how, is Srs2 is removed from the fork under normal circumstances? For what purposes? This, we believe is the next major hurdle in the challenging research involving this protein.

## Materials and Methods

Unless differently stated, all strains are derivatives of E134, and share the following genotype: *ade5-1,lys2::InsEa14,trp1-289,his7-2,leu2-3,112,ura3-52* [38].

Standard yeast media and methods were used. Standard yeast molecular biology techniques were used to create the mutant collection.

**Table 1.**

| Name | Genotype | Source |
| --- | --- | --- |
| 18076 | mat@ ade5-1,lys2::InsEa14,trp1-289,his7-2,leu2-3,112,ura3-52 | [38] |
| 19523 | mat@ ade5-1,lys2::InsEa14,trp1-289,his7-2,leu2-3,112,ura3-52, Pol30 with c' terminal flag with KanMX marker | This study |
| 19787 | matA ade5-1,lys2::InsEa14,trp1-289,his7-2,leu2-3,112,ura3-52, pol30E143K:leu2+flag:kanmx | [29] |
| 19785 | matA ade5-1,lys2::InsEa14,trp1-289,his7-2,leu2-3,112,ura3-52, pol30V180D:leu2+flag:kanmx | [29] |
| 19524 | mat@ ade5-1,lys2::InsEa14,trp1-289,his7-2,leu2-3,112,ura3-52, Elg1::Hyg, Pol30 with c' terminal flag with KanMX marker | [15] |
| 17822 | mat@ ade5-1,lys2::InsEa14,trp1-289,his7-2,leu2-3,112,ura3-52, PCNA V180D:leu | [29] |
| 17823 | mat@ ade5-1,lys2::InsEa14,trp1-289,his7-2,leu2-3,112,ura3-52, PCNA E143K:leu | [29] |
| 19682 | mat@ ade5-1,lys2::InsEa14,trp1-289,his7-2,leu2-3,112,ura3-52,pol30-K164R, K127R:LEU2 | [19] |
| 19808 | matA,ade5-1,lys2::InsEa14,trp1-289,his7-2,leu2-3,112,ura3-52, pol30K164R::leu2 | [19] |
| 20573 | MatA; ade5-1,lys2::InsEa14,trp1-289,his7-2,leu2-3,112,ura3-52, Pol30-k164r;v180d:Leu; | This study |
| 19525 | mat@ ade5-1,lys2::InsEa14,trp1-289,his7-2,leu2-3,112,ura3-52,pol30:E143K,K164R:leu2 | This study |
| 20033 | E134 matA de5-1,lys2::InsEa14,trp1-289,his7-2,leu2-3,112,ura3-52, Srs2::KanMX | [19] |
| 20464 | E134 matA de5-1,lys2::InsEa14,trp1-289,his7-2,leu2-3,112,ura3-52, Srs2::KanMX pol30-V180D: LEU2 | This study |
| 20508 | MAT@ ade5-1,lys2::InsEa14,trp1-289,his7-2,leu2-3,112,ura3-52, Pol30-k164r:LEU, SRS2::KanMX | This study |
| 20463 | E134 matA de5-1,lys2::InsEa14,trp1-289,his7-2,leu2-3,112,ura3-52, Srs2::KanMX Pol30-V180D,K164R:leu2 | This study |
| 20461 | E134 matA de5-1,lys2::InsEa14,trp1-289,his7-2,leu2-3,112,ura3-52, Srs2::KanMX, pol30-K164R, K127R:LEU2 | This study |
| 19678 | mat@ ade5-1,lys2::InsEa14,trp1-289,his7-2,leu2-3,112,ura3-52, pol30-V180D, K164R, K127R: LEU2 - #2 | This study |
| 20462 | E134 matA ade5-1,lys2::InsEa14,trp1-289,his7-2,leu2-3,112,ura3-52, Srs2::KanMX pol30-V180D, K164R, K127R:LEU2 | This study |
| 20609 | E134 A ade5-1,lys2::InsEa14,trp1-289,his7-2,leu2-3,112,ura3-52, rad52::ura3 -1 | This study |
| 20656 | mat@ ade5-1,lys2::InsEa14,trp1-289,his7-2,leu2-3,112,ura3-52, PCNA V180D:leu, rad52::ura3 | This study |
| 20589 | mat@ ade5-1,lys2::InsEa14,trp1-289,his7-2,leu2-3,112,ura3-52, pol30-K164R:leu2, rad52::ura3 | This study |
| 20660 | matA ade5-1,lys2::InsEa14,trp1-289,his7-2,leu2-3,112,ura3-52 Pol30-V180D,K164R:leu2, rad52::ura3 | this study |
| 20708 | Mata; ade5-1,lys2::InsEa14,trp1-289,his7-2,leu2-3,112,ura3-52, Pol30-k164r,k127r::leu Rad52::ura | This study |
| 18045 | mat@ ade5-1,lys2::InsEa14,trp1-289,his7-2,leu2-3,112,ura3-52 elg1 :: hyg | [15] |
| 20545 | Mat@; ade5-1,lys2::InsEa14,trp1-289,his7-2,leu2-3,112,ura3-52, ELG1::Hyg; SRS2:KanMX; | [19] |
| 20544 | Mat@; ade5-1,lys2::InsEa14,trp1-289,his7-2,leu2-3,112,ura3-52, pol30-v180d:leu; ELG1::Hyg; SRS2:KanMX; | This study |
| 20543 | diploid; ade5-1,lys2::InsEa14,trp1-289,his7-2,leu2-3,112,ura3-52, (heterozygous) ELG1::Hyg; (heterozygous) SRS2:KanMX; (heterozygous) Pol30-V180D:leu | This study |
| 20541 | diploid; ade5-1,lys2::InsEa14,trp1-289,his7-2,leu2-3,112,ura3-52, (heterozygous) ELG1::Hyg; (heterozygous) SRS2:KanMX; (heterozygous) Pol30-E143K:leu | This study |

### Chromatin Fractionation Assay

Cells from 50 mL cultures (OD600 < 1.0) were collected by centrifugation, successively washed with ddH2O, PSB (20 mM Tris–Cl pH 7.4, 2 mM EDTA, 100 mM NaCl, 10 mMb-ME), and SB (1 M Sorbitol, 20 mM Tris–Cl pH 7.4), and transferred to a 2 mL Eppendorf tube. Cells were suspended in 1 mL SB, 30ul Zymolase 20T (20 mg/mL in SB) was added, and samples were incubated at 30 °C with rotation until >85% spheroplasts were observed (60–90 min). Spheroplasts were collected by centrifugation (2 K, 5 min, 4 °C), washed twice with SB, and suspended in 500 mL EBX (20mM Tris–Cl pH 7.4, 100 mM NaCl, 0.25% Triton X-100,15 mM-ME + protease/phosphatase inhibitors). TritonX-100 was added to a final concentration of 0.5% to lyse the outer cell membrane, and the samples were kept on ice for 10 min with gentle mixing. The lysate was layered over 1 mL NIB (20 mM Tris–Cl pH 7.4, 100 mM NaCl, 1.2 M sucrose, 15 mM-ME + protease/phosphatase inhibitors) and centrifuged at 12 K RPM for 15 min, at 4 °C. The supernatant (cytoplasm) was discarded. The glassy white nuclear pellet was suspended in 500 uL EBX and Triton X-100 was added to a 1% final concentration to lyse the nuclear membrane. The chromatin and nuclear debris were collected by centrifugation (15 K, 10 min, 4 °C). Chromatin was suspended in 50 uL Tris pH 8.0 for Western blot analysis (Chromatin). To each fraction, an equal volume of 2 × SDS-PAGE loading buffer (60 mM Tris pH 6.8, 2% SDS, 10% glycerol, 0.2%bromophenol blue, 200 mM DTT) was added; samples were incubated at 95 °C for 5 min and were then analyzed by SDS-PAGE and Western blot analyses.

### DNA damage sensitivity assays

Serial 10-fold dilutions of logarithmic yeast cells were spotted on fresh synthetic dextrose (SD)-complete (or SD lacking a specific amino acid to preserve the plasmid) plates with or without different concentrations of methyl methanesulfonate (MMS) (Sigma) and incubated at 30°C for 3 days. MMS plates were freshly prepared, dried in a biological hood, and used the same day.

### Western blot analysis

Protein extracted from the fractionation protocol, either from the chromatin fraction or the WCE were loaded on an acrylamide gel prepared in our lab in 15% acrylamide concentration for PCNA, RPS6 and H3 and on 8% for Srs2. Gels were run for an 90 mintues half in 140V in Bio-Rad western-blot construction. Afterward the gel was taken apart and proteins transferred using Bio-Rad transfer construct for 90 minutes in 400 milliamperes to cellulose membranes. Membrane was incubated for 30 minutes in 1% skim milk for blocking and afterward incubated with primary antibody for over-night (anti-PCNA: Sc65598-santa cruz. anti-Flag:F1804-Sigma Aldrich, anti-H3: ab1791-abcam. Anti-srs2: sc11991-santa cruz. Anti-RPS6:ab40820-abcam). Following morning, membranes are washed 5 times in TTBS, after each wash incubation for 10 min leading to an hour incubation in secondary antibody and afterward another cycle of 5 washes. In the end, using Thermo scientific ECL we expose the membranes in imager600 and capture pictures of the membrane with different exposures.

### Tetrad dissection

Diploid strains are grown overnight (4-5 ml) in YPD, the cells are spined down (2-3 minutes) and resuspended in sporulation medium (SPO) in glass tubes. The culture is incubated at 25°C for 4 days and sporulation is verified under the microscope before continuing to the next step. Spores were treated using beta-glucuronidase and dissected using a micromanipulator. The plates are then grown in 30 degrees for 3 days before photographed and then replica-plated to various plates to test the relevant markers.

## Authors’ contribution

Conceptualization, M.A.-G., B.L. and M.K.; Investigation, M.A.-G., N.K., M.K. (Maxim Kuryachiy); Formal Analysis, M.A.-G., B.L. and M.K.; Writing – Original Draft Preparation, M.A.-G. and M.K.; Writing – Review & Editing, M.A.-G. and M.K.; Supervision, M.K., Funding Acquisition, M.K.

## Acknowledgements

We thank all present and past members of the Kupiec lab for ideas and support.

